# Colorectal cancer progression is potently reduced by a glucose-free, high-protein diet: comparison to anti-EGFR therapy

**DOI:** 10.1101/2021.06.15.448354

**Authors:** Kerstin Skibbe, Ann-Kathrin Brethack, Annika Sünderhauf, Mohab Ragab, Annika Raschdorf, Maren Hicken, Heidi Schlichting, Joyce Preira, Jennifer Brandt, Darko Castven, Bandik Föh, René Pagel, Jens U. Marquardt, Christian Sina, Stefanie Derer

**Author notes:** **Correspondence:** Stefanie Derer, Institute of Nutritional Medicine, University Hospital Schleswig-Holstein, Campus Lübeck, Ratzeburger Allee 160, Tel: +49(0)451/3101 8402; Fax: +49(0)451/3101 8404. these authors contributed equally to this study (Kerstin Skibbe, Ann-Kathrin Brethack). **Grant support:** This work was supported by the German Research Foundation (Research grant DE 1874/1-2 to SD) and an intramural grant from the University of Lübeck (H03-2020 to SD). CS is Fresenius Kabi endowed professor for Nutritional Medicine. **Author contributions:** SD designed the concept of the study and supervised it. KS, AKB, AS, MH, HS, JP, JB, MR, BF, RP and SD performed the experiments and acquired the data. KS, CS and SD analyzed and interpreted the data. SD, AR, MR and AS drafted the article. KS, DC, JUM, CS and SD critically revised the article. All authors read and approved to the final manuscript.

## Abstract

**Background & Aims:** To enable rapid proliferation, colorectal tumor cells up-regulate epidermal growth factor receptor (EGFR) signaling and perform high level of aerobic glycolysis, resulting in substantial lactate release into the tumor microenvironment and impaired anti-tumor immune responses. We hypothesized that an optimized nutritional intervention designed to reduce aerobic glycolysis of tumor cells may boost EGFR-directed antibody (Ab)-based therapy of pre-existing colitis-driven colorectal carcinoma (CRC).

**Methods:** CRC development was induced by azoxymethane (AOM) and dextran sodium sulfate (DSS) administration to C57BL/6 mice. AOM/DSS treated mice were fed a glucose-free, high-protein diet (GFHPD) or an isoenergetic control diet (CD) in the presence or absence of *i*.*p*. injection of PBS, an irrelevant control mIgG2a or an anti-EGFR mIgG2a. *Ex vivo*, health status, tumor load, metabolism, colonic epithelial cell differentiation and immune cell infiltration were studied. Functional validation was performed in murine and human CRC cell lines MC-38 or HT29-MTX.

**Results:** AOM/DSS treated mice on GFHPD displayed reduced systemic glycolysis, resulting in improved tumoral energy homeostasis and diminished tumor load. Comparable but not additive to an anti-EGFR-Ab therapy, GFHPD was accompanied by enhanced tumoral differentiation and decreased colonic PD-L1 and splenic PD-1 immune checkpoint expression, presumably promoting intestinal barrier function and improved anti-tumor immune responses. *In vitro*, glucose-free, high-amino acid culture conditions reduced proliferation but improved differentiation of CRC cells in combination with down-regulation of PD-L1 expression.

**Conclusion:** We here found GFHPD to metabolically reprogram colorectal tumors towards balanced OXPHOS, thereby improving anti-tumor T-cell responses and reducing CRC progression with a similar efficacy as EGFR-directed antibody therapy.

**Synopsis:** Consumption of glucose-free, high protein diet (GFHPD), comparable to EGFR-directed antibody-based therapy, efficaciously dampens colorectal tumor burden but does not have additive effects in combination with anti-EGFR therapy. GFHPD boosts amino acid metabolism, resulting in improved tumor differentiation and anti-tumor immune responses.

**Graphical abstract:** 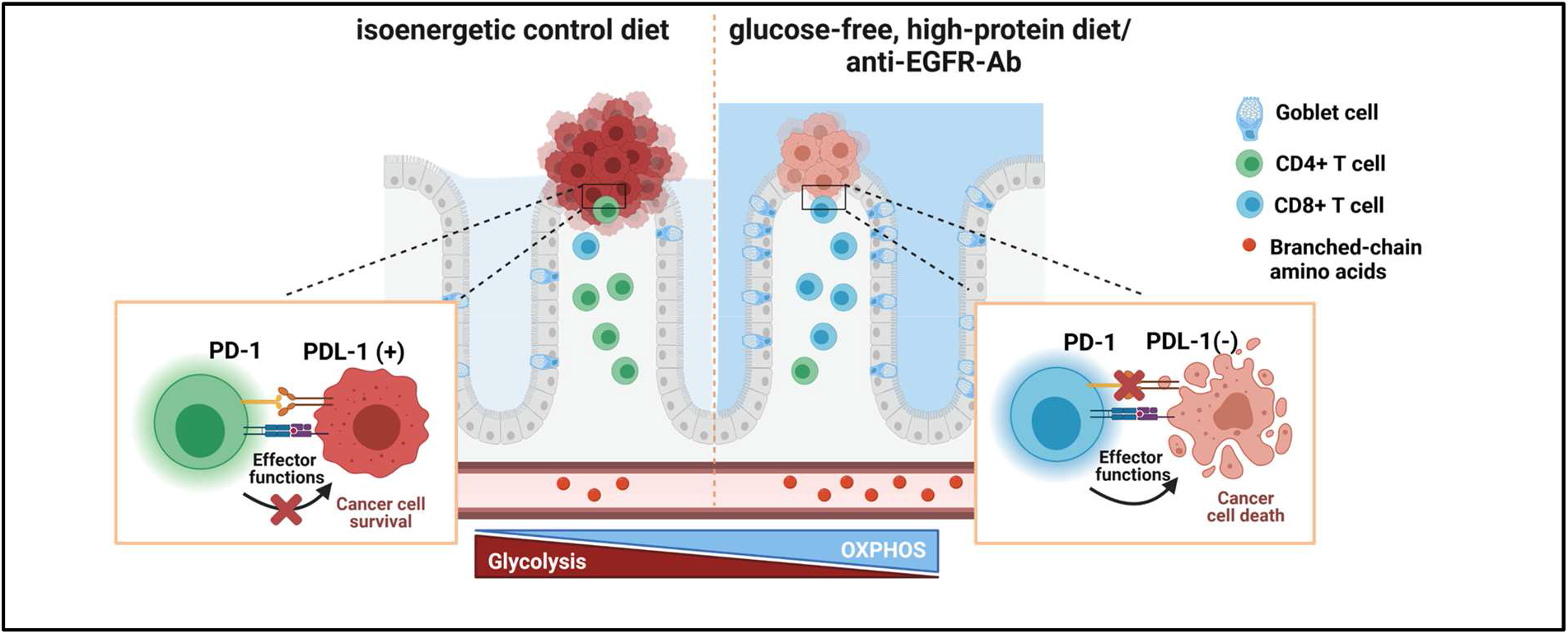

## INTRODUCTION

Colorectal carcinoma (CRC) is one of the most common cancer entities worldwide responsible for more than one million new cases annually [1]. The overall five-year survival rate is about 70% with a recurrence rate of about 5% [2]. It is known that, in addition to genetic predisposition, various aspects of lifestyle, *e*.*g*. physical activity, alcohol consumption, nutrition, and smoking, are important risk factors [3, 4]. Increasing evidence further suggests that chronic inflammation plays a crucial role in the pathogenesis of colorectal cancer [5, 6]. Inflammatory bowel disease (IBD), such as ulcerative colitis (UC) or Crohn’s disease (CD) is an example of the link between inflammation and tumorigenesis, which is a potent risk factor for the development of colorectal cancer [7]. However, the underlying mechanisms remain elusive.

The epidermal growth factor receptor (EGFR) belongs to a family of transmembrane tyrosine kinase receptors. Because of its importance for tumor development and progression, the EGFR has long been considered as a promising target molecule for tumor therapy [8, 9]. Overexpression or mutations of the EGFR are found in a variety of epithelial tumors such as CRC [10]. Therapeutically, EGFR can be targeted by small molecule tyrosine kinase inhibitors (e.g. erlotinib and gefitinib) or by monoclonal antibodies (mAbs) [11]. Currently, three monoclonal antibodies (cetuximab, panitumumab, and necitumumab) have been approved for therapeutic applications, but many more are in clinical development [12]. Approved indications include Kirsten rat sarcoma virus (KRAS) wild type colon cancer for cetuximab and panitumumab, head and neck cancer, where only cetuximab has been approved so far as well as non-small cell lung cancer [13]. In contrast to EGFR blockade in cancer, rectal application of EGF containing enemas to ulcerative colitis patients induced mucosal healing in 10 out of 12 patients, highlighting the crucial role of EGFR signalling cascade in intestinal epithelial regeneration. However, due to the potential activation of pro-oncogenic signalling pathways, this therapeutic strategy was not followed up [14]. In experimental murine colitis models, blockade of EGFR by the tyrosine kinase inhibitor gefitinib [15] or knockout of the EGFR [16] resulted in exacerbated colitis and colitis-driven colorectal carcinoma, indicating a crucial role of EGFR signaling cascades in intestinal tissue homeostasis. However, blockade of EGFR by the tyrosine kinase inhibitor gefitinib in mice displaying established colorectal tumors seemed to decrease polyp sizes, pointing to opposing roles of EGFR signaling in the resolution of intestinal inflammation and in tumor growth [15].

A hallmark of cancer cells is increased glycolysis under normoxic conditions. In contrast to quiescent differentiated cells, tumor cells produce large amounts of lactate instead of CO_2_ during glucose metabolism in the presence of oxygen [17, 18], called “aerobic glycolysis”. This metabolic switch was first described in 1923 by the German biochemist Otto Heinrich Warburg and is also known under the term “Warburg effect” [17]. With a sufficient supply of glucose and oxygen, 85% of glucose in tumor cells enters aerobic glycolysis, a metabolic pathway that enables fast and direct accessible energy [19], while 5% of imported glucose is metabolized *via* the oxidative phosphorylation (OXPHOS) using the mitochondrial respiratory chain [20]. The altered metabolism of a cancer cell is not only used to generate energy in the form of ATP [21], but the intermediates of glycolysis can be diverted into biosynthetic pathways to produce compounds for cell division [22]. Hence, recent studies showed that administration of 2-deoxy-D-glucose (2-DG), a glycolysis inhibitor, reduces the growth of colorectal carcinoma in mice [23].

In triple-negative breast cancer cells, it has been shown that the EGFR signal activates the first step of glycolysis. At the same time, however, this signal also inhibits the last step of glycolysis, resulting in an accumulation of metabolic intermediates such as fructose-1,6-bisphosphate (F1,6BP). The latter enhances its action by directly binding to the EGFR, thereby increasing lactate excretion. Increased lactate production in turn leads to an inhibition of local cytotoxic T cells and a reduced anti-tumoral immune response [24].

To date, only two antibodies against the murine EGFR (mEGFR) are known, the 7A7 [25] and another antibody produced by Imclone [26]. Previous studies show that the so-called 7A7 antibody (7A7-Ab) recognizes the extracellular domain of mEGFR on healthy cells and tumor cells. In addition, 7A7-Ab inhibited EGF-induced signaling cascade of the EGFR in the murine lung carcinoma cell line 3LL-D122 [27, 28]. The inhibition of EGFR-mediated signal transduction induced apoptosis, thereby evoking the initiation of anti-tumor immune responses.

Considering that an acidic tumor microenvironment crucially impairs immune effector cell activation and therefore anti-tumor immune responses [29], we hypothesized that the efficacy of EGFR-targeting antibody-based therapy can be increased by metabolic reprogramming of the tumor itself. Hence, we here investigated the therapeutic efficacy of an optimized nutritional intervention designed to inhibit increased aerobic glycolysis of tumor cells in the presence or absence of an EGFR-directed antibody-based therapy in a mouse model of colitis-driven CRC. Therefore, we applied the sporadic azoxymethane (AOM)/dextran sodium sulfate (DSS) triggered mouse model of colitis-driven colorectal cancer [30, 31]. AOM is a genotoxic carcinogen that causes DNA damage and tumorigenesis and specifically induces the development of colorectal cancer in rodents [32]. The combination of AOM with the inflammatory drug DSS is widely used in colitis-driven colon carcinogenesis in mouse models [33].

## RESULTS

### Anti-EGFR Ab therapy or glucose-free, high-protein nutritional intervention efficaciously prevent CRC progression

To study the effect of nutritional intervention in the presence or absence of anti-EGFR Ab treatment on the progression of pre-existing CRC in mice, we first induced sporadic colitis-driven CRC in C57BL/6 mice until day 70 by intraperitoneal (*i*.*p*.) injection of 10 mg/kg/body weight (BW) AOM at day 0 and a second *i*.*p*. injection of 5 mg/kg/BW AOM at day 21. Colitis was induced by the application of 2% (w/v) DSS *via* the drinking water between day 7 and 14 and was boosted by a second application of 1% (w/v) DSS *via* the drinking water between day 42 and 45. C57BL/6 control mice underwent *i*.*p* injections of 1x PBS instead of AOM and remained on normal drinking water **(Fig. 1a)**. Before *in vivo* application, we studied the binding of utilized monoclonal murine IgG2a antibody 7A7 (anti-EGFR Ab) to murine recombinant EGFR protein by ELISA experiments and determined a significant, concentration-dependent binding, while the control antibody CD19-mIgG2a (ctrl. Ab) did not show any binding capacity to murine EGFR **(Fig. 1b)**. During the first 70 days, the disease activity index (DAI), combining weight loss, stool consistency, and rectal bleeding, was monitored every two days and indicated successful colitis induction in AOM/DSS treated mice in comparison to PBS control mice **(Fig. 1c)**. On day 70, indicator mice were sedated and endoscopically examined before sacrificing. As presented in figure 1d, AOM/DSS treated mice displayed colorectal tumor development, while no tumors were endoscopically detected in PBS treated mice at day 70 **(Fig. 1d)**. As expected from human CRC studies, increased *Egfr* mRNA expression was detected in colorectal tumors from AOM/DSS treated mice in comparison to normal tissue **(Fig. 1e)**.

**Figure 1.**
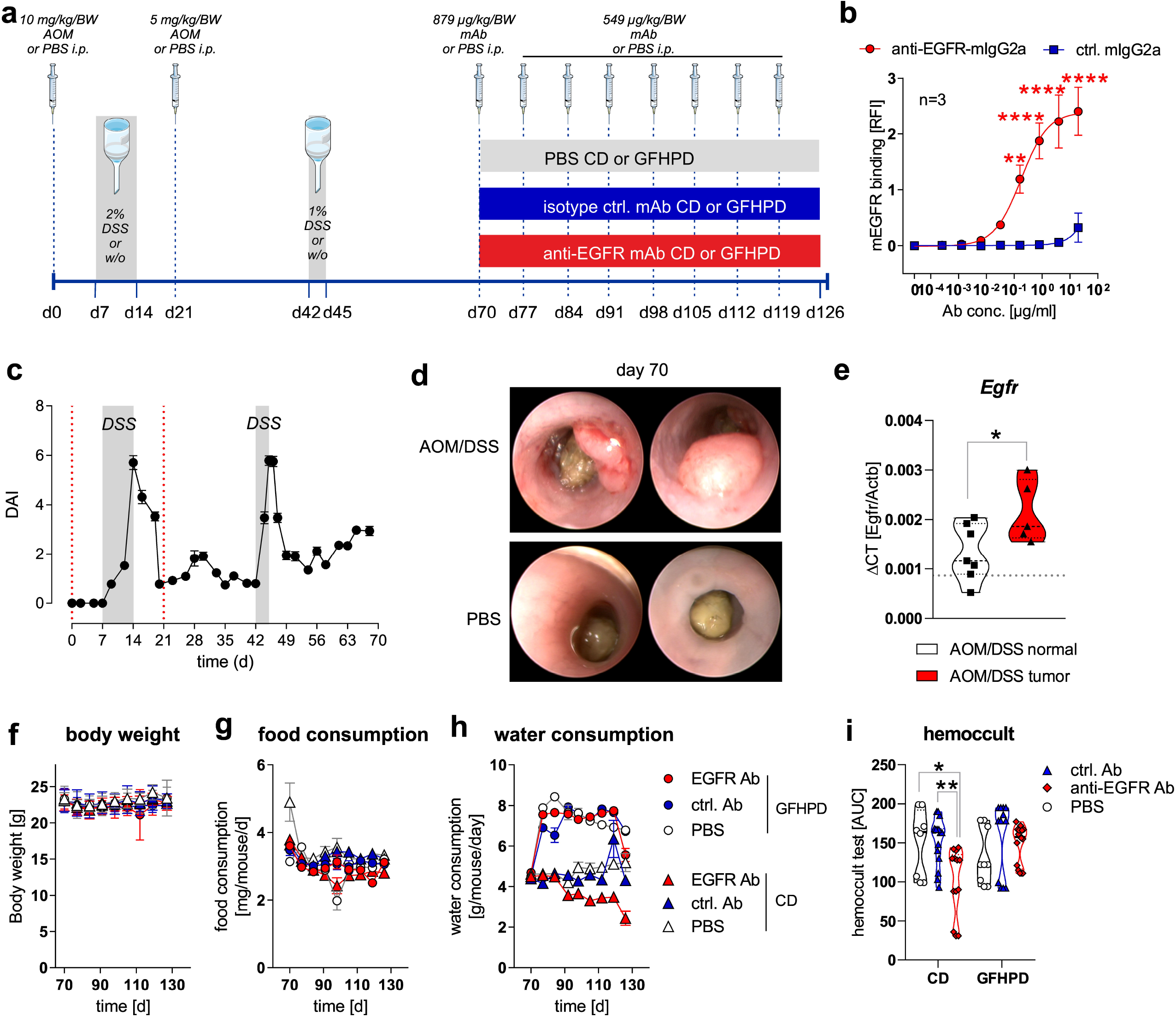
**(a)** Schematic overview of induction of colitis-driven colorectal carcinomas by AOM/DSS treatment in female C57BL/6J mice (n=88) between day 0 and 70. Control mice received PBS instead of AOM/DSS (n=19) between day 0 and 70. At day 70, indicator mice (n=9 AOM/DSS treated; n=8 PBS treated) were sacrificed and tissues were sampled for further analyses. Remaining AOM/DSS treated mice were further subdivided at day 70 into the two different dietary intervention groups receiving control diet (CD; n=39) or glucose-free, high-protein diet (GFHPD; n=40) until day 126. Dietary interventions were performed in the presence or absence of weekly intraperitoneal (*i*.*p*.) application of PBS (n=9 CD; n=9 GFHPD), anti-EGFR Ab (n=15 CD; n=16 GFHPD) or a ctrl. antibody (n=15 CD; n=15 GFHPD) between day 70 and 126. PBS-treated control mice were fed either CD (n=5) or GFHPD (n=6) between day 70 and 126. AOM = azoxymethane; DSS = dextran sodium sulfate; anti-EGFR-mAb = 7A7-mIgG2a; isotype ctrl. mAb = irrelevant mIgG2a; mAb = monoclonal Ab. **(b)** Concentration-dependent binding of anti-EGFR-mIgG2a or ctrl.-mIgG2a to recombinant murine EGFR was analyzed by ELISA experiments. Mean ± SEM of three independent experiments is presented. **(c)** Disease activity index (DAI) of AOM/DSS treated mice (n=88) from day 0 to 70. **(d)** Representative images of endoscopy analyses of AOM/DSS or PBS treated indicator mice at day 70. **(e)** *Egfr* mRNA expression was quantified by qPCR and related to *Actb* mRNA expression in normal or tumoral colonic tissues collected from AOM/DSS treated mice at day 70. Grey dotted line indicates median values pf PBS treated control mice at day 70. **(f-h)** Body weight, food consumption and water consumption were routinely monitored between day 70 and 126. **(i)** Fecal occult blood was determined every second day using a hemoccult test. Results are presented as the area under the curve (AUC) from analyzed mice. * *p* ≤ 0.05, ** *p* ≤ 0.01. **** *p* ≤ 0.0001.

From day 70 on, AOM/DSS treated or PBS treated mice were separated into two subgroups, with one group receiving control diet (CD; 20% protein/casein, 5% fat, 53% carbohydrates, 15% fibre; **tab. 1**) and one receiving an isoenergetic glucose-free, high-protein diet (GFHPD; 60% protein/casein, 11% fat, 20% fibre; **tab. 1**) as a therapeutic regimen. Furthermore, AOM/DSS treated mice either in the CD or the GFHPD groups additionally received an initial dose of 0.879 mg/kg/BW of anti-EGFR Ab, ctrl. Ab or PBS at day 70 and a weekly dose of 0.549 mg/kg/BW from day 77 until day 126 by *i*.*p*. injection **(Fig. 1a)**. Regularly, body weight, food and water consumption as well as fecal blood were determined to monitor the health status of all analyzed mice. While no differences were observed between nutritional intervention and antibody treatment groups regarding body weight **(Fig. 1f)** and food consumption **(Fig. 1g)**, all mice on the GFHD, irrespective of antibody treatment, displayed significantly increased water consumption in comparison to CD mice **(Fig. 1h)**. Of note, hemoccult test revealed anti-EGFR Ab treatment to significantly decrease frequency of fecal blood in CD but not in GFHPD fed AOM/DSS treated mice compared to PBS or ctrl. Ab treated mice **(Fig. 1i)**.

As expected, in AOM/DSS treated mice on CD, anti-EGFR Ab treatment markedly lowered frequency of large tumors with a score of 5 (ctrl. Ab = 50% of tumors with tumor score 5, anti-EGFR Ab = 22.2% of tumors with tumor score of 5) compared to ctrl. Ab. Similarly, the application of anti-EGFR Ab to AOM/DSS treated mice fed a GFHPD significantly reduced the presence of large tumors with a tumor score of 5 (27.3%) when compared to ctrl. Ab application (57.1%), resulting in a higher percentage of tumors with a tumor score of 4 (27.3%), while no tumors with a tumor score of 4 were detected in GFHPD fed mice treated with the ctrl. Ab. Compared to CD, GFHPD did not alter the presence of large colonic tumors of tumor score 5 in AOM/DSS and ctrl. Ab treated mice. However, consumption of a GFHPD by AOM/DSS and ctrl. Ab treated mice lowered the presence of colonic tumors with a score between 1-4, therefore increasing the percentage of tumor-free mice to 43% in comparison to 20% tumor-free mice in the CD group **(Fig. 2a+b)**. Results received from tumor scoring analyses were reflected by endoscopic examination of the colon that indicated a significantly reduced murine endoscopic index of colitis severity (MEICS) in AOM/DSS and ctrl. Ab treated mice fed a GFHPD (mean ± SD; 1.8 ± 1.2) in comparison to CD (mean ± SD; 4.1 ± 1.7). While anti-EGFR Ab treatment resulted in a trend towards a reduced MEICS score (mean ± SD; 2.9 ± 2.2) in AOM/DSS mice fed a CD in comparison to ctrl. Ab treatment, the MEICS score was slightly increased but without statistical significance by anti-EGFR Ab treatment in AOM/DSS mice fed a GFHPD diet (mean ± SD; 3.2 ± 1.4) **(Fig. 2c)**.

**Figure 2.**
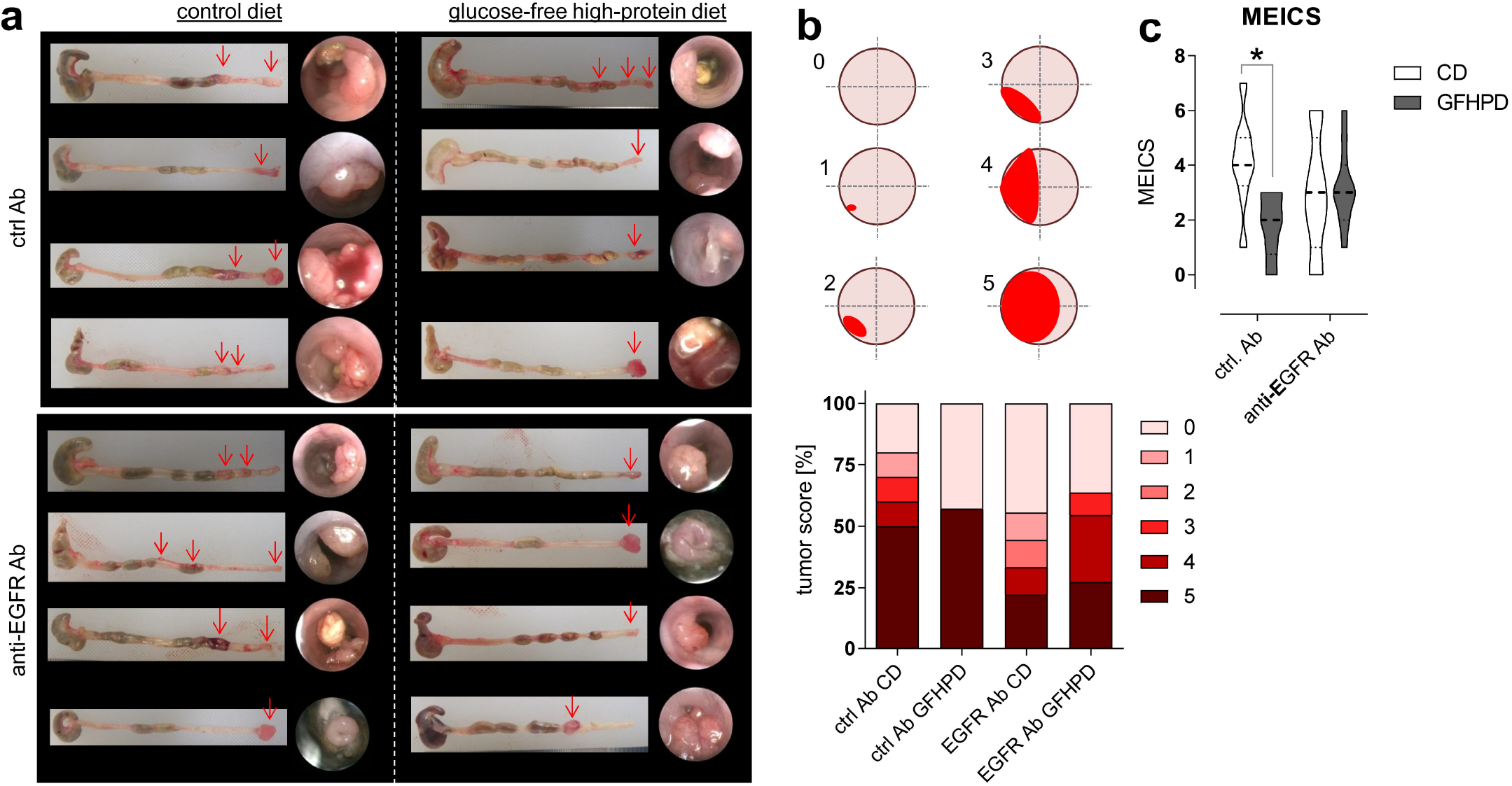
**(a)** Representative images (n=4 per indicated group) from colons collected from AOM/DSS treated mice and respective endoscopy analyses at day 126. **(b)** Colorectal tumors were scored between 0 and 5 as described in [60]. Percentages for each tumor score was calculated per treatment group. **(c)** Murine endoscopic index of colitis severity (MEICS) was determined accordingly to [60]. * *p* ≤ 0.05.

### A glucose-free, high-protein diet increases amino acid metabolism and decreases lactate and insulin production

To investigate the effect of GFHPD consumption on systemic and colonic metabolism in AOM/DSS treated mice, we analyzed metabolic markers in serum and colonic biopsy samples collected at day 126 from ctrl. Ab or anti-EGFR Ab treated mice. Here, in ctrl. Ab treated mice L-lactate serum levels were found to be significantly reduced in GFHPD fed mice in comparison to CD fed mice, while no alterations were detected after anti-EGFR Ab therapy **(Fig. 3a)**. Of note, comparable glucose serum levels were determined in all analyzed groups **(Fig. 3b)**, while GFHPD lowered insulin levels without reaching statistical significance in ctrl. Ab treated mice (mean ± SD, 2.8 ± 6.3 µiU/ml) when compared to CD fed mice (mean ± SD, 6.9 ± 6.1 µiU/ml; **Fig. 3c**). In line with insulin serum levels, C-peptide serum levels were significantly reduced in ctrl. Ab treated mice fed a GFHPD **(Fig. 3d)**. Furthermore, anti-EGFR Ab therapy in CD fed mice did not alter insulin or C-peptide serum levels in comparison to ctrl. Ab treatment, while GFHPD fed mice showed similarly reduced insulin but increased C-peptide serum levels after anti-EGFR Ab therapy (mean ± SD, 0.7 ± 3.1 µiU/ml) compared to ctrl. Ab treated mice **(Fig. 3c+d)**. As the expected physiological reaction to high protein consumption, branched-chain amino acid and L-alanine serum levels were significantly enhanced in ctrl. Ab treated mice fed a GFHPD compared to a CD, while no alterations were observed in all other treatment groups **(Fig. 3e+f)**. Due to the significantly elevated water consumption and serum amino acid load in GFHPD fed mice, we studied kidney morphologies in order to exclude a kidney failure. Here, no microscopic signs of kidney injury could be detected by HE staining analysis in any analyzed group **(Fig. 3g)**.

**Figure 3.**
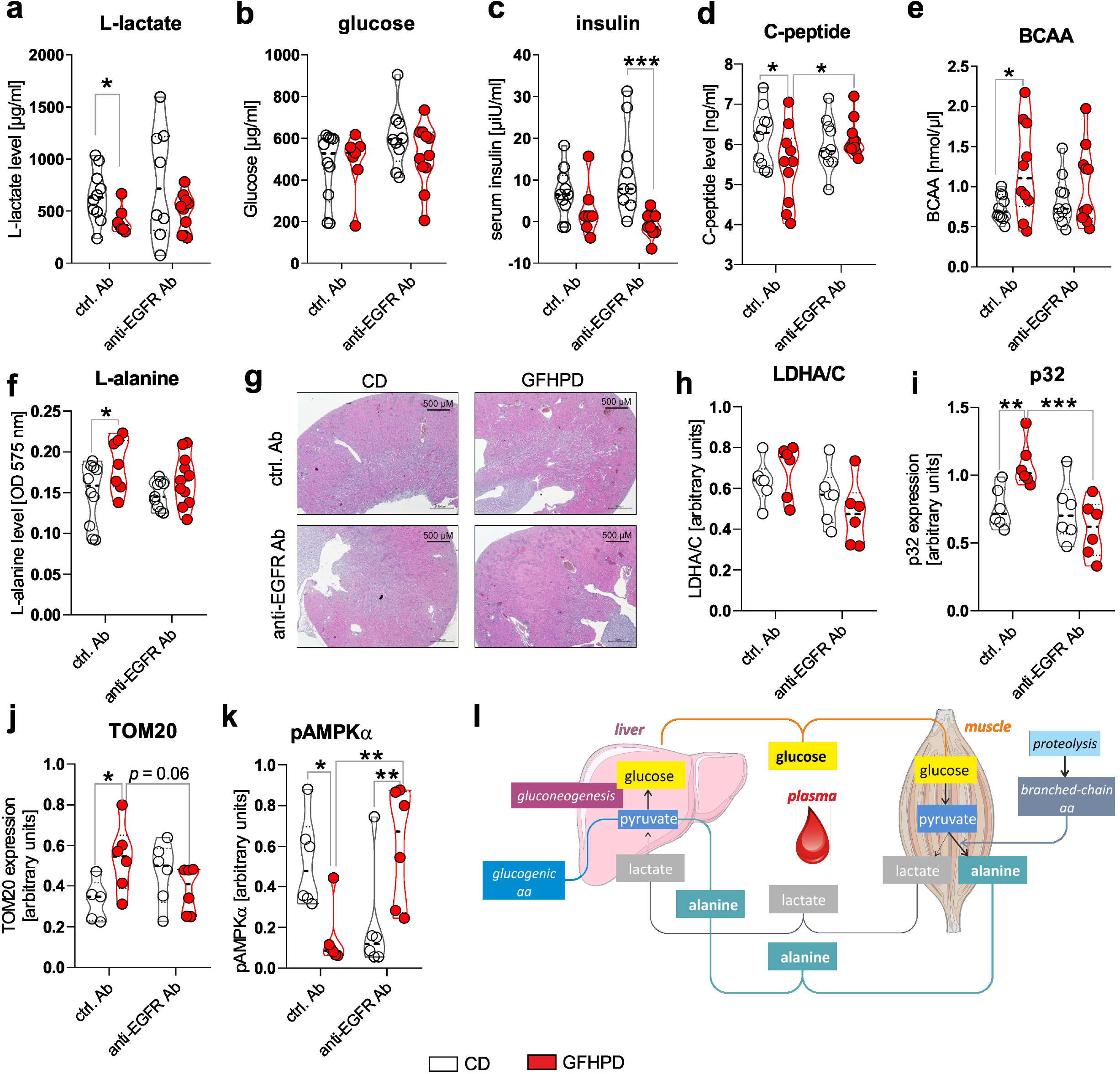
**(a-f)** Serum samples collected from indicated treatment groups at day 126 were analyzed with regard to (a) L-lactate, (b) D-glucose, (c) insulin, (d) C-peptide, (e) branched-chain amino acids (BCAA) or (f) L-alanine. ctrl. Ab (n=10 CD; n=7-10 GFHPD), anti-EGFR Ab (n=8-10 CD; n=9-11 GFHPD) **(g)** Representative images of HE-stained kidney tissue collected at day 126 from indicated treatment groups of AOM/DSS tumor mice. Original magnification, 2.5x. **(h-k)** Protein expression of (h) LDHA/C, (i) p32, (j) TOM20, (k) phosphorylated AMPKα or α-Tubulin was determined by western bot experiments utilizing biopsy samples collected from colorectal tumors (n=6 per group) of indicated treatment groups at day 126. Densitometric analysis was performed using ImageJ software and data were normalized to α-Tubulin values. **(l)** Schematic overview of systemic amino acid metabolism. * *p* ≤ 0.05, ** *p* ≤ 0.01. *** *p* ≤ 0.001.

In the next set of experiments, we investigated the impact of increased systemic amino acid and decreased glucose metabolism on colonic metabolism. Therefore, we performed western blot experiments utilizing colonic biopsy samples from AOM/DSS and ctrl. Ab or anti-EGFR Ab treated mice fed a CD or a GFHPD. Despite the decreased systemic glucose metabolism in GFHPD mice, no alterations in colonic lactate dehydrogenase a/c (LDHA/C) protein expression could be observed between all four groups **(Fig. 3h)**. Of note, expression of mitochondrial markers such as P32 **(Fig. 3i)** and translocase of outer mitochondrial membrane 20 (TOM20; **Fig. 3j**) was significantly increased in AOM/DSS and ctrl. Ab treated mice fed a GFHPD when compared to CD, revealing the GFHPD to boost mitochondrial mass and therefore mitochondrial function. Furthermore, enhanced expression of mitochondrial markers P32 and TOM20 in ctrl. Ab treated mice were significantly down-regulated in anti-EGFR Ab treated mice under GFHPD. In GFHPD fed mice, up-regulation of colonic mitochondrial mass was accompanied by a reduction of AMPKα phosphorylation and hence energy deficiency under ctrl. Ab treatment that was elevated to the level of CD fed mice under anti-EGFR Ab treatment **(Fig. 3k)**. Metabolic analyses of AOM/DSS mice indicate that a GFHP nutritional intervention in contrast to a control diet reduces systemic glucose metabolism and increases amino acid metabolism to maintain constant glucose serum level, resulting in improved colonic mitochondrial OXPHOS activity that is abrogated under anti-EGFR Ab treatment plus GFHPD **(Fig. 3l)**.

### Goblet cell differentiation is enhanced by GFHP diet or anti-EGFR Ab therapy in AOM/DSS treated mice

Cell proliferation and differentiation processes in the colon are driven by caspase-1 mediated metabolic switches between aerobic glycolysis and balanced mitochondrial OXPHOS activity, respectively, and need to be tightly controlled to maintain proper colonic barrier function [34, 35]. Due to findings from metabolic analyses presented in figure 3, we further investigated mRNA expression levels of *Caspase-1* as well as colonic epithelial cell differentiation markers *Lgr5, Hes1, Atoh1, Spdef1*, and *Klf4* **(Fig. 4a)**. Notably, no differences in colon length as a marker for colitis activity could be detected between all four analyzed treatment groups **(Fig. 4b)**, while *Caspase-1* mRNA was reduced (*p* = 0.051) in ctrl. Ab treated and GFHPD fed mice when compared to CD mice **(Fig. 4c)**, pointing to decreased colonic cell proliferation under GFHPD. As depicted in figure 4d-g, we did not detect any differences between analyzed mice in mRNA expression of the intestinal stem cell marker *Lgr5* **(Fig. 4d)**, the enterocyte marker *Hes1* **(Fig. 4e)**, the secretory cell precursor marker *Atoh1* **(Fig. 4f)**, and the goblet cell precursor marker *Spdef1* **(Fig. 4g)**. In line with reduced mRNA expression of the intestinal cell proliferation mediator *Caspase-1*, we found a significantly increased mRNA expression of the goblet cell differentiation markers *Klf4* and *Muc2* in GFHPD fed *versus* CD fed AOM/DSS mice that were treated with a ctrl. Ab. Furthermore, *Klf4* mRNA expression was also significantly increased in anti-EGFR Ab *versus* ctrl. Ab treated mice fed a CD, while *Muc2* expression was significantly reduced in anti-EGFR Ab *versus* ctrl. Ab treated mice fed a GFHPD **(Fig. 4h+i)**.

**Figure 4.**
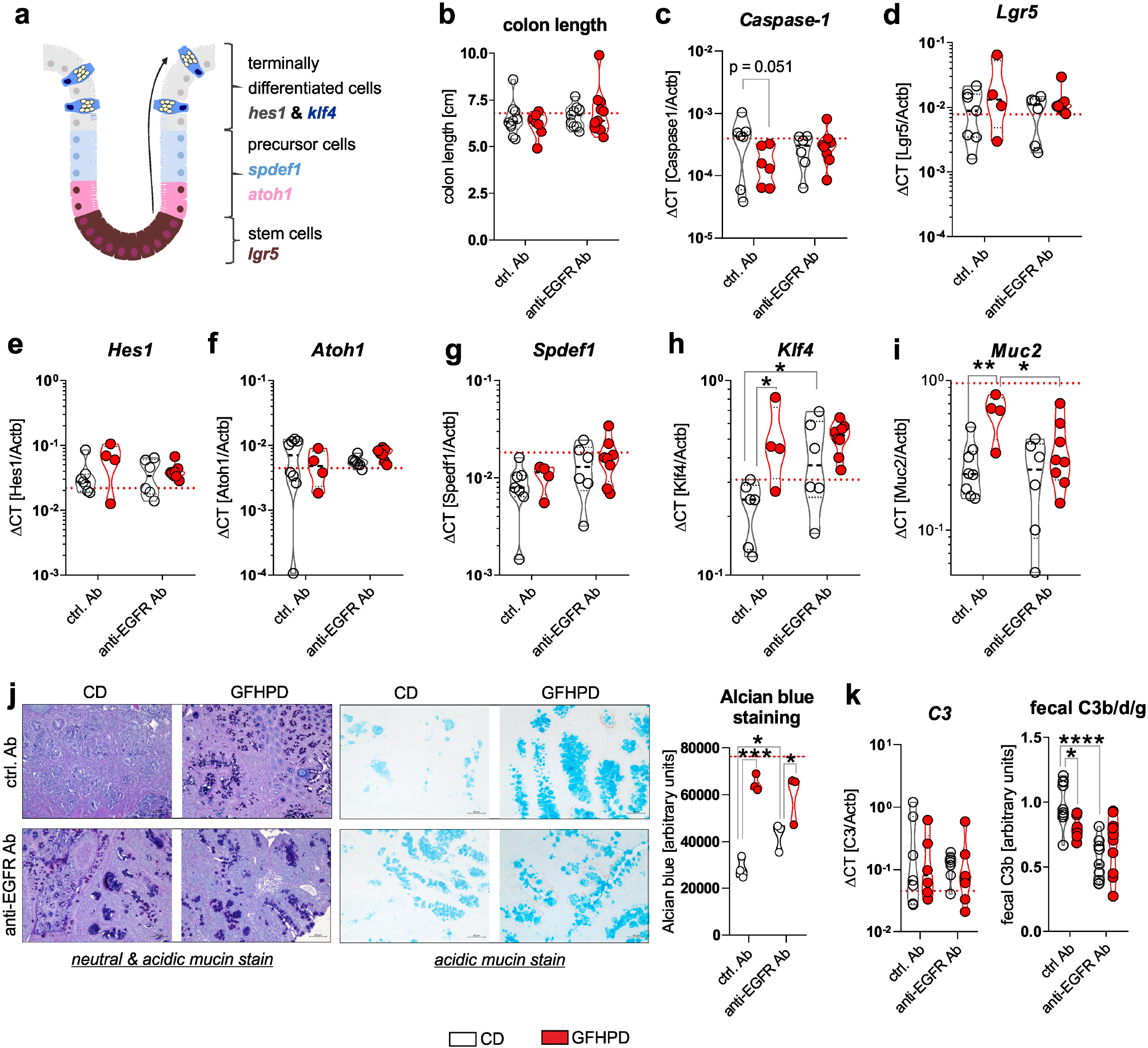
**(a)** A model for intestinal epithelial cell differentiation in the colonic crypt. **(b)** Whole colon length was determined at day 126. **(c-i)** MRNA expression levels of (c) *Caspase-1*, (d) *Lgr5*, (e) *Hes1*, (f) *Atoh1*, (g) Spdef1, (h) *Klf4* or (i) *Muc2* were quantified by qPCR in colonic biopsy samples collected from indicated treatment groups. Ctrl. Ab (n=6-8 CD; n=4-6 GFHPD), anti-EGFR Ab (n=6-7 CD; n=8-9 GFHPD). **(j)** Representative histochemical PAS/Alcian (left panel) or Alcian blue (right panel) stainings of PFA-fixed, paraffin-embedded tumor tissue biopsies from indicated treatment groups. Original magnification, 10x. **(k)** Colonic *C3* mRNA expression was quantified by qPCR experiments (left panel). Ctrl. Ab (n=7 CD; n=6 GFHPD), anti-EGFR Ab (n=7 CD; n=8 GFHPD). Fecal C3b/d/g protein levels were measured by western blot experiments utilizing fecal samples collected from indicated treatment groups of AOM/DSS mice (right panel). Densitometric analysis was performed using the ImageJ software. Ctrl. Ab (n=10 CD; n=7 GFHPD), anti-EGFR Ab (n=9 CD; n=10 GFHPD). Red dotted horizontal lines represent respective median values of PBS-treated control mice at day 126. * *p* ≤ 0.05, ** *p* ≤ 0.01. **** *p* ≤ 0.0001.

For validation of qPCR data, we performed histochemical staining experiments. Here, increased neutral & acidic mucin expression was determined by PAS/Alcian blue **(Fig. 4j, left panel)** or Alcian blue **(Fig. 4j, right panel)** staining of colonic tumors in either GFHPD fed mice (ctrl. Ab or anti-EGFR Ab treated) or in anti-EGFR Ab treated mice fed CD **(Fig. 4j)**. As a functional consequence of enhanced mucus production, we analyzed fecal C3 cleavage products that have been shown previously to be triggered by gram-negative mucosa-associated bacteria *via* Toll-like receptor 4 (TLR4) signaling in intestinal epithelial cells and hence to constitute a fecal marker of intestinal barrier integrity [36]. While colonic *C3* mRNA expression did not differ between analyzed groups **(Fig. 4k, left panel)**, fecal C3b/d/g protein levels were significantly decreased in GFHPD fed mice treated with a ctrl. Ab or in CD fed mice treated with an anti-EGFR Ab in comparison to ctrl. Ab treated mice fed a CD **(Fig. 4k, right panel)**. Fecal C3 cleavage products were not further reduced in GFHPD fed mice plus anti-EGFR Ab therapy. Together, GFHPD consumption enhances colorectal goblet cell differentiation in AOM/DSS and ctrl. Ab treated mice to a comparable extent as anti-EGFR Ab therapy, resulting in an improved intestinal barrier function.

### Increased amino acid metabolism enhances goblet cell differentiation in colorectal tumor cells *in vitro*

To mechanistically validate the described *ex vivo* results, *in vitro* experiments were performed utilizing the murine colorectal carcinoma cell line MC-38 and the human colorectal carcinoma cell line HT29-MTX. MC-38 cells were incubated for 72 hours in culture media containing increasing concentrations of glucose (0 – 25 mM) in the presence of 1% non-essential amino acids (NEAA), 10% NEAA, or 20% NEAA to study cell proliferation, differentiation, and lactate production. In concordance with *in vivo* data, MC-38 cell viability was increased in a glucose-concentration-dependent manner that was further elevated in the presence of increasing NEAA concentrations **(Fig. 5a)**. Of note, MC-38 cells that were cultivated in a glucose-free medium enriched with NEAA (1%, 10% and 20%) displayed significantly decreased cell proliferation in comparison to the standard culture medium (25 mM glucose, 1% NEAA) **(Fig. 5b)**. Decreased cell proliferation of MC-38 cells under glucose-free plus 20% NEAA conditions after 48 hours of cultivation was also reflected by decreased mRNA and protein expression of proliferation driving Caspase-1 **(Fig. 5c)** and reduced activation of the main proliferation pathway *via* AKT **(Fig. 5d)**. Consistent with these findings, incubation of HT29-MTX cells under glucose-free plus 10% NEAA conditions also demonstrated significantly reduced cell proliferation in comparison to glucose-containing plus 1% NEAA conditions **(Fig. 5e)**. As we observed decreased lactate serum levels in AOM/DSS and ctrl. Ab treated mice fed a GFHPD *versus* CD, we measured lactate release of MC-38 and HT29-MTX cells. Similar to cell proliferation results, lactate release by MC-38 cells increased in a glucose-concentration-dependent manner after 72 hours of incubation. While the highest lactate release was detected in MC-38 cells cultivated for 72 hours in 25 mM glucose plus 1% NEAA containing medium, a lower one was achieved by the addition of 20% NEAA to the culture medium **(Fig. 5f)**. Likewise, under glucose-free culture conditions, lactate release by MC-38 cells was lowest in the presence of 20% NEAA **(Fig. 5g)**. These results were validated in HT29-MTX cells that displayed significantly decreased lactate release per cell in glucose-free plus 10% NEAA culture conditions in comparison to 25 mM glucose plus 1% NEAA culture conditions **(Fig. 5h)**. To investigate whether decreased cell proliferation of CRC cell lines in glucose-free plus high amino acid conditions results in enhanced goblet cell differentiation, we studied goblet cell marker levels in MC-38 and HT29-MTX cells. Whereas mRNA expression of the goblet cell marker *Klf4* was not significantly altered by metabolic challenges in MC-38 cells, *Muc2* mRNA expression and Muc2 secretion were significantly increased under glucose-free plus 20% NEAA culture conditions when compared to 25 mM glucose plus 1% NEAA culture conditions after 48 hours of incubation **(Fig. 5i)**. As expected, a significantly elevated Muc5AC release was observed in HT29-MTX cells incubated for 72 hours in glucose-free plus 10% NEAA culture conditions in comparison to cultivation under 25 mM glucose plus 1% NEAA **(Fig. 5j)**. In summary, a reduction of glucose metabolism and enhanced amino acid metabolism improves goblet cell differentiation in colorectal carcinoma cells.

**Figure 5.**
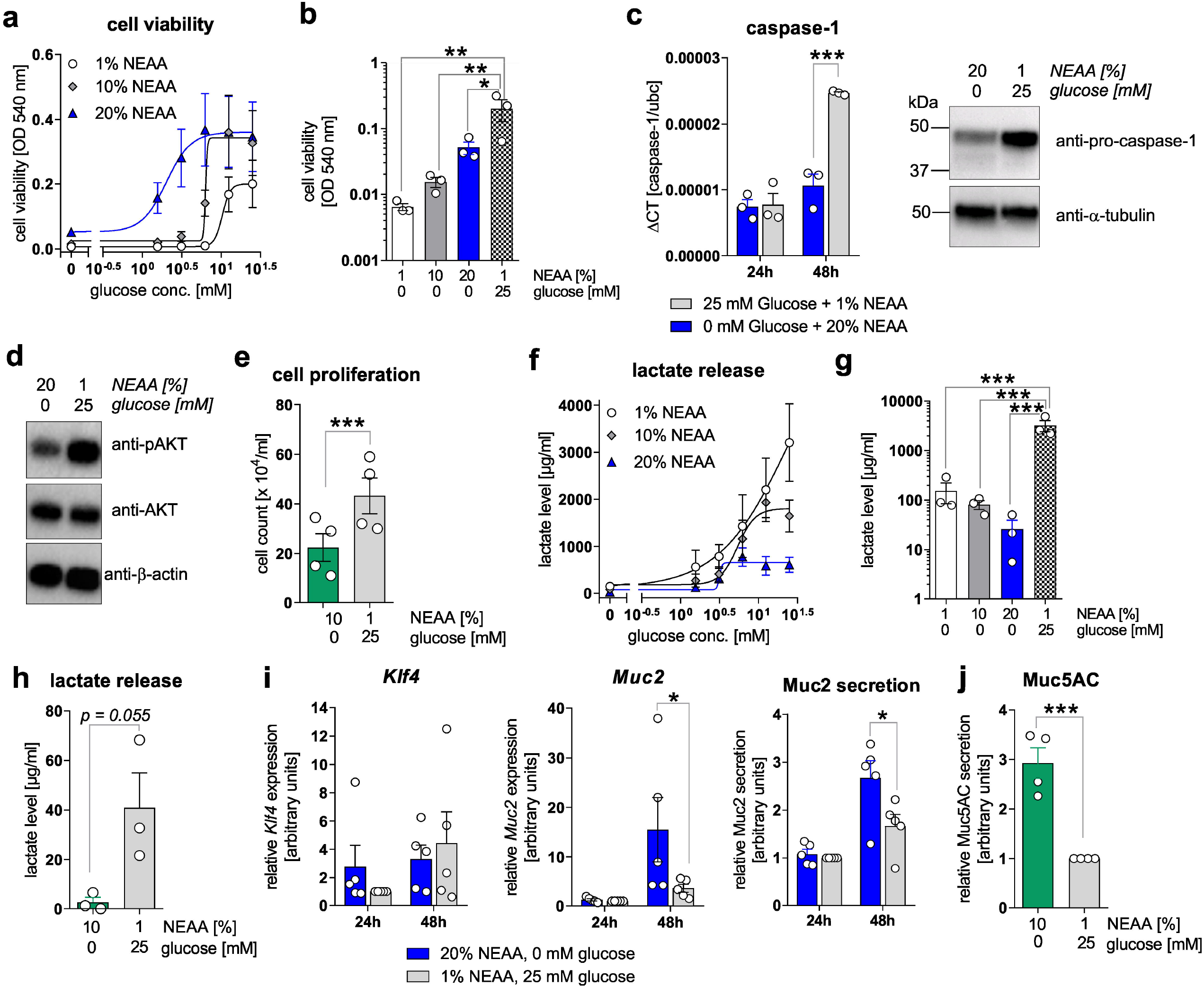
**(a+b)** MC-38 cells were incubated in culture media containing increasing concentrations of D-glucose (0-25 mM) in the presence of 1%, 10% or 20% NEAA. After 72 hours, cell viability was determined by neutral-red cytotoxicity assay. **(c)** MC-38 cells were incubated in standard culture medium (25 mM glucose, 1% NEAA) or in glucose-free plus 20% NEAA medium for 24 or 48 hours. *Caspase-1* mRNA expression was quantified by qPCR experiments and related to *Ubc* mRNA expression (left panel; n=3 independent experiments). Caspase-1 protein expression after 48 hours of incubation was analyzed by western blot experiments. Alpha-Tubulin expression served as a loading control (right panel; representative images of n=3 independent experiments). **(d)** Protein levels of phosphorylated AKT and AKT were determined after 48 hours of incubation of MC-38 cells under indicated culture conditions by western blot experiments. β-actin expression served as a loading control. Representative images of n=3 independent experiments. **(e)** HT29-MTX cells were incubated for 72 hours under indicated culture conditions and cell count was determined. Mean ± SEM of four independent experiments is presented. **(f+g)** MC-38 cells were incubated under indicated culture conditions. After 72 hours, L-lactate levels were quantified in cell culture supernatants. **(h)** L-lactate release by HT29-MTX cells after 72 hours of incubation under indicated culture conditions. L-lactate concentrations were related to respective cell count presented in (e). Mean ± SEM of three independent experiments is presented. **(i)** MC-38 cells were incubated in standard culture medium (25 mM glucose, 1% NEAA) or in glucose-free plus 20% NEAA medium for 24 or 48 hours. *Klf4, Muc2* or *Ubc* mRNA expression was quantified by qPCR experiments (n=3 independent experiments). ΔCT values were calculated by normalization of *Klf4* or *Muc2* to *Ubc* and related to values received from standard culture conditions at 24h. Muc2 secretion was analyzed after 72 hours of incubation in cell culture supernatants from respective cells by ELISA experiments. Data are related to cell viability determined by neutral-red cytotoxicity assay and presented as mean ± SEM. **(j)** Muc5AC release by HT29-MTX cells was measured after 72 hours of incubation and related to respective cell counts. * *p* ≤ 0.05, ** *p* ≤ 0.01. *** *p* ≤ 0.001.

### The metabolic switch from glucose to amino acid metabolism as well as anti-EGFR Ab treatment decreases PD-L1 expression in CRC cells and colonic CD4^+^ T-cell count

Metabolic active tumors create a tumor environment which is characterized by a high tumoral lactate release resulting from high aerobic glycolysis activity that has been demonstrated to inhibit anti-tumor immune responses of T-cells and macrophages [18]. Hence, we analyzed mRNA expression of leukocyte markers in colonic biopsy samples of analyzed mice **(Fig. 6a)**. While no differences in the expression of the pan-leukocyte marker *Cd45* were detected, the T-cell specific *Cd4* marker but not *Cd3e* or *Cd8a* was significantly down-regulated in GFHPD *versus* CD fed mice treated with a ctrl. Ab. Furthermore, anti-EGFR Ab therapy did not alter *Cd3e* or *Cd4* expression in CD fed mice but did significantly upregulate both transcripts in GFHPD fed mice **(Fig. 6b+c)**. No differences in mRNA expression were detected for markers specific for natural killer (NK) cells (*Cd69*; **Fig. 6a**), plasma cells (*Bcma*; **Fig. 6a**), macrophages (*F4/80*; **Fig. 6a**), or granulocytes (*Ly6g*; **Fig. 6a**). Based on the strong inhibitory effect of GFHPD on CD4^+^ T-cell infiltration into the colonic compartment in AOM/DSS mice, we further quantified colonic expression of the immune checkpoints *Pd-1* and *Ctla4* as well as their respective ligands *Pd-l1* and *Cd80* or *Cd86* **(Fig. 6a)**. Here, *Pd-1, Ctla4, Cd80* or *Cd86* mRNA expression was comparable between analyzed groups, while a trend towards down-regulation without statistical significance of *Pd-l1* expression in GFHPD (mean ± SD, 0.00034 ± 0.00025) *versus* CD (mean ± SD, 0.001 ± 0.00034) fed mice treated with a ctrl. Ab was observed. Of note, anti-EGFR Ab treatment under CD displayed a significantly lower *Pd-l1* expression when compared to anti-EGFR Ab treatment under GFHPD **(Fig. 6d)**.

**Figure 6.**
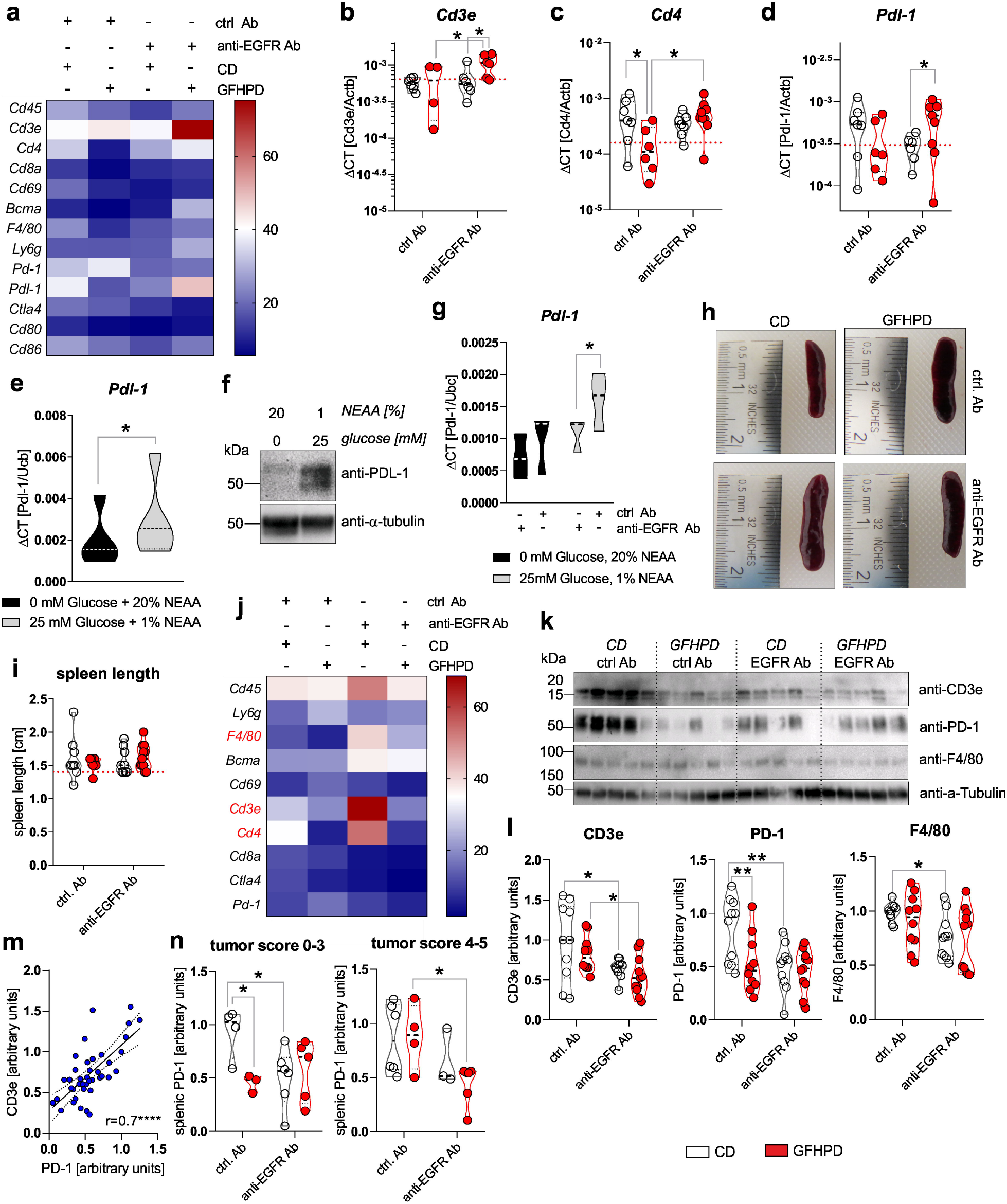
**(a-d)** QPCR analyses were performed to quantify mRNA expression level of Cd45, Cd3e, Cd4, Cd8a, Cd69, Bcma, F4/80, Ly6g, Pd-1, PD-L1, Ctla4, Cd80 or Cd86 in colonic tissues collected from indicated treatment groups of AOM/DSS mice. Results are based on ΔCT calculation including relation to β-actin mRNA expression. Heatmap was generated based on normalized values (%) per transcript. The mean value per goup is presented. n=6-7 ctrl. Ab & CD; n=6-7 anti-EGFR Ab & CD; n=4-6 ctrl. Ab & GFHPD; n=6-9 anti-EGFR Ab & GFHPD. Red dotted horizontal lines represent respective median values of PBS-treated control mice at day 126. **(e+f)** MC-38 cells were incubated in standard culture medium (25 mM glucose, 1% NEAA) or in glucose-free plus 20% NEAA medium for 48 hours. (e) *PD-L1* mRNA expression was quantified by qPCR experiments and related to *Ubc* mRNA expression (n=5 independent experiments), while (f) PD-L1 protein expression was analyzed by western blot experiments. Alpha-Tubulin expression served as a loading control (representative images of n=5 independent experiments). **(g)** MC-38 cells were incubated in standard culture medium (25 mM glucose, 1% NEAA) or in glucose-free plus 20% NEAA medium in the presence or absence of either the anti-EGFR Ab 7A7-mIgG2a or the irrelevant control Ab CD19-mIgG2a for 48 hours. *PD-L1* mRNA expression was quantified by qPCR experiments and related to *Ubc* mRNA expression (n=3 independent experiments). **(h)** Representative images from spleens collected from AOM/DSS treated mice at day 126. **(i)** Representative spleen lengths were calculated for indicated treatment groups. n=9 ctrl. Ab & CD; n=9 anti-EGFR Ab & CD; n=6 ctrl. Ab & GFHPD; n=11 anti-EGFR Ab & GFHPD. Red dotted horizontal line represents median values of PBS-treated control mice at day 126. **(j)** QPCR analyses were performed to quantify mRNA expression levels of Cd45, Ly6g, F4/80, Bcma, Cd69, Cd3e, Cd4, Cd8a, Ctla4 and Pd-1 in splenic tissues collected from indicated treatment groups of AOM/DSS mice. Results are based on ΔCT calculation including relation to β-actin mRNA expression. Heatmap was generated based on normalized values (%) per transcript. The mean value per group is presented. n=10 ctrl. Ab & CD; n=10 anti-EGFR Ab & CD; n=10 ctrl. Ab & GFHPD; n=11 anti-EGFR Ab & GFHPD. **(k)** Splenic CD3e, PD-1 and F4/80 protein expression levels were determined by western blot experiments. Alpha-Tubulin expression served as a loading control. One representative western blot analysis is presented. **(l)** Densitometric analyses were performed using ImageJ software. n=10 ctrl. Ab & CD; n=10 anti-EGFR Ab & CD; n=10 ctrl. Ab & GFHPD; n=11 anti-EGFR Ab & GFHPD. **(m)** Splenic PD-1 protein expression was related to splenic CD3e protein expression. **(n)** Splenic PD-1 protein expression is displayed in relation to colorectal tumor scores (smaller tumor 0-3; larger tumor 4-5). * *p* ≤ 0.05, ** *p* ≤ 0.01. **** *p* ≤ 0.0001.

To functionally validate results received from *ex vivo* experiments, PD-L1 mRNA as well as protein expression was determined in murine colorectal carcinoma cells MC-38. As depicted in figure 6e+f, incubation of MC-38 cells in glucose-free plus 20% NEAA culture conditions for 48 hours significantly reduced PD-L1 mRNA and protein expression in comparison to 25 mM glucose plus 1% NEAA culture conditions **(Fig. 6e+f)**. Additionally, we incubated MC-38 cells in the presence of either an anti-EGFR-mIgG2a Ab or a ctrl. Ab under glucose-free plus 20% NEAA or 25 mM glucose plus 1% NEAA culture conditions for 48 hours. Of note, anti-EGFR Ab treatment of MC-38 cells induced a significant downregulation of *Pd-l1* exclusively under glucose-containing plus 1% NEAA culture conditions, while no alterations were detected under glucose-free and 20% NEAA culture conditions **(Fig. 6g)**.

Together, these results demonstrate a significant up-regulation of PD-L1 in highly glycolytic CRC cells that can be prevented by GFHPD or anti-EGFR Ab application.

### PD-1 expression on splenic T cells is reduced under GFHP diet or anti-EGFR Ab treatment in AOM/DSS treated mice

Next, we investigated the effect of GFHPD consumption as well as anti-EGFR Ab treatment on leukocyte marker expression in the splenic compartment that is crucially involved in anti-tumor immune responses [37]. Macroscopically, no differences were found between analyzed the groups regarding the size of the spleens **(Fig. 6h+i)**. Next, qPCR experiments were performed to identify potentially regulated subtypes of splenic leukocytes **(Fig. 6j)**. No differences in mRNA expression levels were detected for *Cd45, Ly6g, Bcma, Cd69, Cd8a, Ctla4*, and *Pd-1*. However, anti-EGFR Ab therapy in CD fed mice was accompanied by the upregulation of *F4/80, Cd3e,* and *Cd4*. In contrast, GFHPD consumption in ctrl. Ab treated mice strongly reduced splenic *Cd3e* and *Cd4* expression as already observed in the colonic compartment **(Fig. 6j)**. Next, western blot experiments were performed with protein extracts from respective spleens collected from analyzed mice **(Fig. 6k)**. In contrast to qPCR results, anti-EGFR Ab treatment was accompanied by significantly lower CD3ε and PD-1 protein expression levels under CD when compared to ctrl. Ab treatment group. Furthermore, PD-1 but not CD3ε expression was strongly reduced in ctrl. Ab treated mice fed a GFHP diet when compared to those mice fed a CD. However, anti-EGFR Ab therapy of mice under GFHPD displayed a significant down-regulation of splenic CD3ε expression in comparison to ctrl. Ab therapy. Notably, GFHPD consumption did not alter expression of the macrophage marker F4/80, while a significant reduction was detected under anti-EGFR Ab therapy in CD fed mice when compared to ctrl. Ab therapy **(Fig. 6l)**. As splenic PD-1 expression highly correlated with splenic CD3e expression **(Fig. 6m)**, we subdivided PD-1 protein expression levels into mice harboring colorectal tumors with a score of 0-3 or 4-5. In mice harboring smaller colorectal tumors (score of 0-3), splenic PD-1 expression was significantly reduced under GFHP diet (ctrl. Ab therapy) *vs*. CD or under anti-EGFR Ab therapy plus CD. In contrast, mice harboring larger colorectal tumors (score 4-5), displayed significantly reduced splenic PD-1 expression exclusively after anti-EGFR Ab treatment under GFHPD, while no differences were observed under CD **(Fig. 6n)**.

In summary, GFHPD consumption efficiently reduced PD-1 and PDL-1 immune-checkpoint protein expression to a comparable extent as anti-EGFR Ab therapy, suggesting both therapeutic regimens to boost anti-tumoral T-cell responses.

## DISCUSSION

In this study, we identified a glucose-free, high-protein diet (GFHPD) to induce a metabolic reprogramming from glucose to amino acid metabolism, thereby reducing CRC burden with similar efficacy as EGFR-directed antibody therapy in an experimental mouse model of colitis-driven CRC. Furthermore, therapeutic application of GFHPD or anti-EGFR Ab to mice harboring established CRC tumors significantly improved tumoral energy homeostasis and hence cell differentiation reflected by increased goblet cell function. Of note, GFHPD but not anti-EGFR Ab therapy significantly increased tumoral expression of mitochondrial markers such as p32 and TOM20. In line with these findings, our group recently published caspase-1 mediated cleavage of the mitochondria marker p32 to prevent its mitochondrial translocation and hence to boost aerobic glycolysis in tumor cells, a process that is amplified in high-grade CRC tumors [35]. Furthermore, p32-driven OXPHOS highly drives colonic goblet cell differentiation in normal large intestine [34]. According to studies demonstrating mucus production to be strongly decreased in CRC [38-40], heterozygous expression of the most prominent single nucleotide polymorphism (SNP) in p32 *rs56014026* was characterized to maintain quiescent metabolism and goblet cell differentiation of CRC cells and to be expressed in low-grade colorectal carcinomas in patients [41]. Hence, GFHPD associated increase of tumoral p32 expression in combination with decreased energy deficiency and improved goblet cell differentiation points to metabolic reprogramming of colorectal carcinomas towards balanced OXPHOS, thereby preventing CRC tumor progression. In contrast, anti-EGFR antibody therapy directly inhibited growth factor-mediated tumor cell proliferation without boosting mitochondrial mass. The concept of dietary carbohydrate reduction in tumor therapy constitutes an old concept that is gaining increased acceptance today [42]. One of the first studies directly demonstrating that the isoenergetic replacement of some carbohydrate portions by proteins reduces skin or mammary tumor formation in mice was published by Tannenbaum in 1945 [43]. These data were further validated by Ho *et al*. who further decreased dietary carbohydrate content to 8% combined with an isoenergetic replacement by increased protein fraction. In this study, murine squamous cell carcinoma cell line SCCVII or human colorectal carcinoma cell line HCT-116 xenografts displayed diminished tumor growth in male but not female mice fed a low-carbohydrate and high-protein diet [44].

Besides, enhancement of CRC goblet cell differentiation, both therapeutic regimens, GFHPD or anti-EGFR antibody therapy, strongly dampened colonic PD-L1 and splenic T-cell specific PD-1 immune checkpoint expression, presumably promoting improved anti-tumor immune responses. In detail, ctrl. Ab therapy under GFHPD intervention compared to CD did not alter colonic *Cd3e* or *Cd8a* expression, while *Cd4* and *Pd-l1* were downregulated. Hence, one may hypothesize that GFHPD consumption mainly reduces expansion/infiltration of colonic CD4^+^ but not of CD8^+^ T-cells. This is in line with findings demonstrating high tumoral CD8^+^ but not CD3^+^ cell density to be associated with a lower risk of recurrence in CRC patients [45, 46]. CD8^+^ T cells, especially CD8^+^ cytotoxic T cells, display one of the major immune cell types for tumor cell destruction that is a high-energy demanding immune cell, mainly performing high glycolysis and OXPHOS activities as well as glutaminolysis [47]. In the splenic compartment, we observed a strong down-regulation of CD3ε and PD-1 expression under ctrl. Ab therapy plus GFHPD as well as under anti-EGFR antibody therapy (CD and GFHPD). Due to data revealing PD-1 to be prominently expressed on CD4^+^ tumor-infiltrating lymphocytes (TILs) and to a lesser extent on CD8^+^ TILs [48], we could speculate that CD4^+^ T cells highly depend on glycolysis, while CD8^+^ T cells are able to better compensate low glycemic diets. Furthermore, due to the strong inhibition of anti-tumor T-cell responses *via* binding of tumor cell-expressed PD-L1 to T-cell expressed PD-1 [48, 49], decreased PD-1 expression on circulating splenic T cells in combination with decreased colonic PD-L1 expression in the present study may result in improved anti-tumor T-cell responses in CRC mice treated either with GFHPD or with an anti-EGFR antibody. Recent publications also unraveled EGFR to be expressed on CD4^+^ T cells and that the EGFR-specific tyrosine kinase inhibitor erlotinib inhibits CD4^+^ T-cell proliferation *in vitro*, while a CD4^+^ T-cell specific deletion of *Egfr* in mice was also accompanied by a loss of T-cell activation and proliferation *in vivo [50]*. Hence, anti-EGFR antibody therapy applied to CRC mice in the present study may reduce splenic CD4^+^ T-cell load *via* EGFR blockade on CD4^+^ T cells as well as by induction of Fc-mediated effector cell cytotoxicity against CD4^+^ T cells [51].

One additional major advantage of a glucose-free, high protein nutritional intervention as a single therapeutic regimen in CRC patients may be the independence of tumoral mutation profiles or EGFR cell surface expression levels. In contrast to nutritional intervention, the efficacy of anti-EGFR antibody therapy strongly depends on the tumoral EGFR expression level [51, 52] as well as on the absence of activating mutations of the RAS protein family members [53-55]. Hence, one may hypothesize that the combination of GFHPD and anti-EGFR Ab therapy may be beneficial in low EGFR expressing CRC tumors, while the single GFHPD application may be efficacious in the therapy of RAS mutated CRC tumors.

Notably, in the present study, we applied a diet to mice with pre-existing CRC tumors that was completely deprived of glucose and instead enriched by the protein source casein to achieve a reduction of glycolysis and pentose-phosphate pathway and an increase of mitochondrial OXPHOS. Unexpectedly, consumption of this glucose-free, high-protein diet for more than two months was not accompanied by weight loss, a reduction of blood glucose level, or altered behavior of mice. These data highlight the hypothesis that mice are able to compensate for nutritional glucose deprivation and to use a protein source to maintain blood glucose levels. However, diets high in protein fraction are known to enhance renal function and may affect kidney health. Indeed, we observed a significant increase in water consumption by mice fed a GFHP diet but without any morphological impact on analyzed kidneys. Previously, consumption of a protein-rich diet was accompanied by increased renal blood flow and elevated intraglomerular pressure, resulting in enhanced glomerular filtration rates [56-58]. In the present study, casein, a protein derived from cow’s milk, served as the major source of energy. One disadvantage of this chosen GFHPD may be the transfer to nutritional intervention studies in humans. Interestingly, besides high fibre or low alcohol intake, diets rich in dairy proteins have been associated with a decreased risk of CRC development in humans [4, 59].

Together, we here provide a novel therapeutic strategy to reduce CRC burden by applying a glucose-free, high-protein containing diet as a therapeutic regimen that systematically shifts glucose metabolism to amino acid metabolism, thereby promoting tumoral mitochondria function. Furthermore, single application of a GFHPD to CRC bearing mice displayed comparable therapeutic efficacy as EGFR-directed antibody-based therapy, while the combination of both strategies did not result in an additive effect.

## METHODS

### Animal experiments

All animal experiments were approved by the ethics committee, Schleswig-Holstein, Germany (V 242 – 27664/2018 (64-5/17)) and were performed in two independent experimental rounds. Mice were maintained at the University of Lübeck under specific pathogen-free conditions at a regular 12-hour light–dark cycle with access to food and water *ad libitum*. Procedures involving animals and their care were conducted in accordance with national and international laws and regulations. Glucose free high protein diet (GFHPD) and isoenergetic control diet (CD) were purchased from Ssniff (Soest, Germany). Compositions of corresponding diets are specified in **table 1**. Female C57BL/6 mice (n=107) were ordered at an age of 7-8 weeks from Charles River (Wilmington, Massachusetts, US) and were left to acclimatize on a standard chow diet (Altromin #1324, Lage, Germany). At the age of 10 weeks (day 0), one group (n=88) initially received 10 mg/kg/BW AOM *via* intraperitoneal (*i*.*p*.) injection, while the second control group (n=19) received *i*.*p*. injection of 1x PBS. Experimental chronic colitis was induced in mice on day 7 by administration of 2% (w/v) dextran sodium sulfate (DSS; Batch DB001-27, molecular mass 40 kDa; TdB Consultancy, Uppsala, Sweden) dissolved in the drinking water for 7 d, followed by normal drinking water *ad libitum*. A second *i*.*p*. injection of 5 mg/kg/BW AOM or 1x PBS was performed on day 21. On day 42, mice either received a second administration of 1% (w/v) DSS *via* the drinking water or were left untreated until day 45. On day 70, indicator mice (n=9 AOM/DSS treated; n=8 PBS treated) mice were euthanized and high-resolution mouse video endoscopy was applied (Hopkins Optik 64019BA; Aida Vet; Karl Storz, Tuttlingen, Germany) to obtain the murine endoscopic index of colitis severity (MEICS) [60]. Remaining mice were randomly distributed on day 70 into GFHPD (n=40) and isoenergetic CD (n=39) receiving groups until day 126. Mice were kept on the corresponding diet on an average of 56 days before sampling. In AOM/DSS treated mice (d0-70), the two dietary intervention groups were subdivided into following groups of weekly *i*.*p*. applications: 1x PBS (n=9 CD; n=9 GFHPD), anti-EGFR Ab (7A7-mIgG2a [25, 27, 28]; initial dose of 879 µg/kg/BW at day 70, weekly dose of 549 µg/kg/BW; n=15 CD; n=16 GFHPD) or an irrelevant ctrl. antibody (CD19-mIgG2a; initial dose of 879 µg/kg/BW at day 70, weekly dose of 549 µg/kg/BW; n=15 CD; n=15 GFHPD). PBS-treated control mice (d0-70) were fed either CD (n=5) or GFHPD (n=6) between day 70 and 126. Food and water consumption and body weight were monitored weekly. The modified disease activity index (DAI) was assessed routinely as a combined score of weight loss (0 = none, 1 = 1-5%, 2 = 5-10%, 3 = 10-15%, 4 = 15-20%), stool consistency (0 = solid, 2 = soft, 3 = diarrhea) and rectal bleeding, determined by a fecal occult blood test, (0 = no blood, 2 = positive, 4 = gross blood). Animals that lost more than 20% of their initial weight were euthanized. At the end of the experiment, colonoscopy was performed in sedated animals utilizing high-resolution mouse video endoscopy (Hopkins Optik 64019BA, Aida Vet) and tumor scores were determined according to [60]. Afterwards, mice were sacrificed and clinical parameters were assessed.

**Table 1:** Chow composition as provided by the manufacturer.

| Diet composition | Isoenergetic control diet | Glucose free high protein diet |
| --- | --- | --- |
| Casein | 20% | 60% |
| Brewer`s yeast | - | 2% |
| Corn starch | 28% | - |
| Maltodextrin | 14.5% | - |
| Sucrose | 10% | - |
| Cellulose powder | 15% | 19.5% |
| L-Cystine | 0.3% | 0.3% |
| Vitamin premixture | 1% | 1% |
| Minerals & trace elements | 6% | 6% |
| Choline chloride | 0.2% | 0.2% |
| Soybean oil | 5% | 11% |
| Crude protein | 17.7% | 53.4% |
| Crude fat | 5.1% | 11.4% |
| Crude fibre | 15.8% | 20.3% |
| Crude ash | 5.4% | 5.9 % |
| Starch | 26.9% | 0.1% |
| Sugar | 9.9% | - |
| NfE | 51.7% | 1.9% |
| Physiological fuel value | 13.6 MJ/kg | 13.6 MJ/kg |
| Protein | 22 kcal% | 66 kcal% |
| Fat | 14 kcal% | 32 kcal% |
| Carbohydrates | 64 kcal% | 2 kcal% |

2 NfE: nitrogen-free extract

### Histology and microscopy analyses

Histochemical staining in paraformaldehyde (PFA)-fixed and paraffin-embedded tissue biopsies was performed according to standard protocols. After deparaffinization, rehydration, endogenous peroxidase blockage and antigen retrieval, tissue slides were stained with hematoxylin-eosin (HE), periodic acid-Schiff (PAS) or Alcian blue solutions. Images were obtained and analyzed on an Axio Scope.A1 microscope (Zeiss, Oberkochen, Germany) utilizing the ZEN imaging software (Zeiss).

### Metabolic analyses in serum samples

Murine whole blood samples were collected, placed on ice for 30 minutes and centrifuged at 500 x g and 4°C for 5 minutes. Serum samples were transferred to new vials and stored at -80°C. Serum levels of L-lactate (Megazyme, Wicklow, Ireland), glucose (Analyticon^®^ Biotechnologies AG, Lichtenfels, Germany), insulin (Thermo Fisher Scientific Inc., Waltham, Massachusetts, USA), C-peptide (RayBiotech Life, Inc., Peachtree Corners, GA, US), branched-chain amino acids (BCAA; Sigma Aldrich, St. Louis, US) or L-alanine (AAT Bioquest, Inc., Sunnyvale, CA, US) were measured in diluted serum samples according to manufacturer’s instructions. Optical densities were measured on a SpectraMax iD3 microplate reader (Molecular Devices, San José, California, US).

### Cell culture

The murine colorectal carcinoma cell line MC-38, derived from C57BL/6 mice, (Kerafast Inc., Boston, Massachusetts, US) and the human colorectal carcinoma cell line HT29-MTX-E12 (Sigma Aldrich, St. Louis, US) were kept in DMEM medium supplemented with 1% non-essential amino acids (NEAA). All cell culture media were supplemented with 10% (v/v) heat-inactivated FCS, 100 U/ml penicillin, and 100 mg/ml streptomycin. Cells were incubated at 37 °C and 5% CO_2_ in a humidified incubator. Cells were cultivated up to a maximum of 20 passages and confirmed to be negative for mycoplasma contamination every three months and when freshly thawed. For D-glucose and NEAA titration experiments, cells were seeded in glucose-free DMEM medium supplemented with 10% FCS, 100 U/ml penicillin and 100 mg/ml streptomycin and D-glucose and/or NEAA solutions were added at indicated concentrations.

### RNA extraction, cDNA synthesis and quantitative PCR

RNA was extracted with the innuPREP RNA mini kit (Analytik Jena AG, Jena, Germany) and transcribed to cDNA (RevertAid H Minus reverse transcriptase; Thermo Fisher Scientific) using the T Gradient thermocycler (Whatman Biometra, Göttingen, Germany). Quantitative PCR (qPCR) was performed on the StepOne real-time system (ThermoFisher Scientific) applying Perfecta SYBR Green Supermix (ThermoFisher Scientific). Following cycling conditions were applied: initial denaturation at 95°C for 5 min; 40 cycles of denaturation at 95°C for 45 sec, annealing at appropriate temperature (55°C) for 30 sec and elongation for 30 sec at 72°C. Melting curve profiles were produced and data were analyzed following the 2^−ΔCt^ algorithm by normalization to *β-actin* or *ubc*. Primer sequences are listed in **table 2**.

**Table 2:**
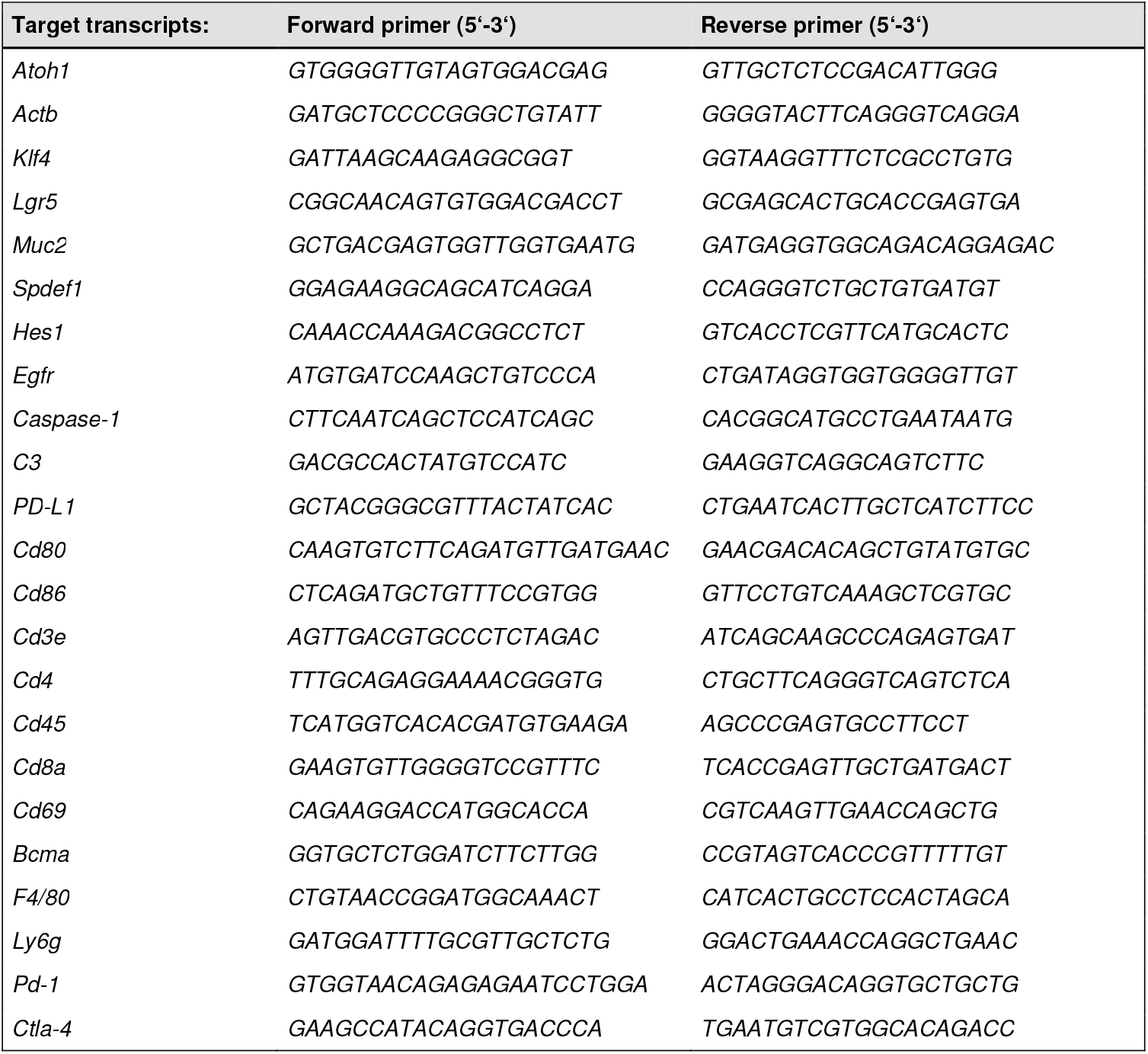
Information on applied primer pairs.

### Sodium dodecyl sulfate polyacrylamide gel electrophoresis (SDS-PAGE) and immunoblotting

SDS-PAGE and immunoblotting was performed according to standard protocols. In short, whole-protein extracts from homogenized tissue samples or cells were prepared by cell lysis in denaturing lysis buffer containing 1% (w/v) SDS, 10 mM Tris (pH 7.4), phosphatase II, phosphatase III and protease inhibitor (Sigma Aldrich). For extraction of proteins from fecal samples, 50 mg of feces were resuspended in 1 ml 0.5% (v/v) Tween, 0.05% (w/v) sodium azide in PBS and centrifuged at 12,000 x *g* for 15 min at 4°C. Supernatant was taken off, protease inhibitor cocktail (Sigma-Aldrich, St. Louis, MO, USA) was added, samples were centrifuged as stated above and supernatant was taken for further analysis. Protein extracts were separated by denaturing SDS-PAGE (Bio-Rad Laboratories, Hercules, California, US) under reducing conditions and transferred onto polyvinylidene difluoride membranes. After blocking, membranes were probed with specific primary antibodies followed by respective HRP-conjugated secondary antibodies. To determine similar transfer and equal loading, membranes were stripped and reprobed with an appropriate housekeeper. Proteins of interest were visualized on a ChemiDocTM XRS^+^ Imaging System (Bio-Rad Laboratories). Applied antibodies are listed in **table 3**. Densitometric analyses were performed using the software ImageJ version 1.53e (National Institutes of Health, Bethesda, MD, USA).

**Table 3:**
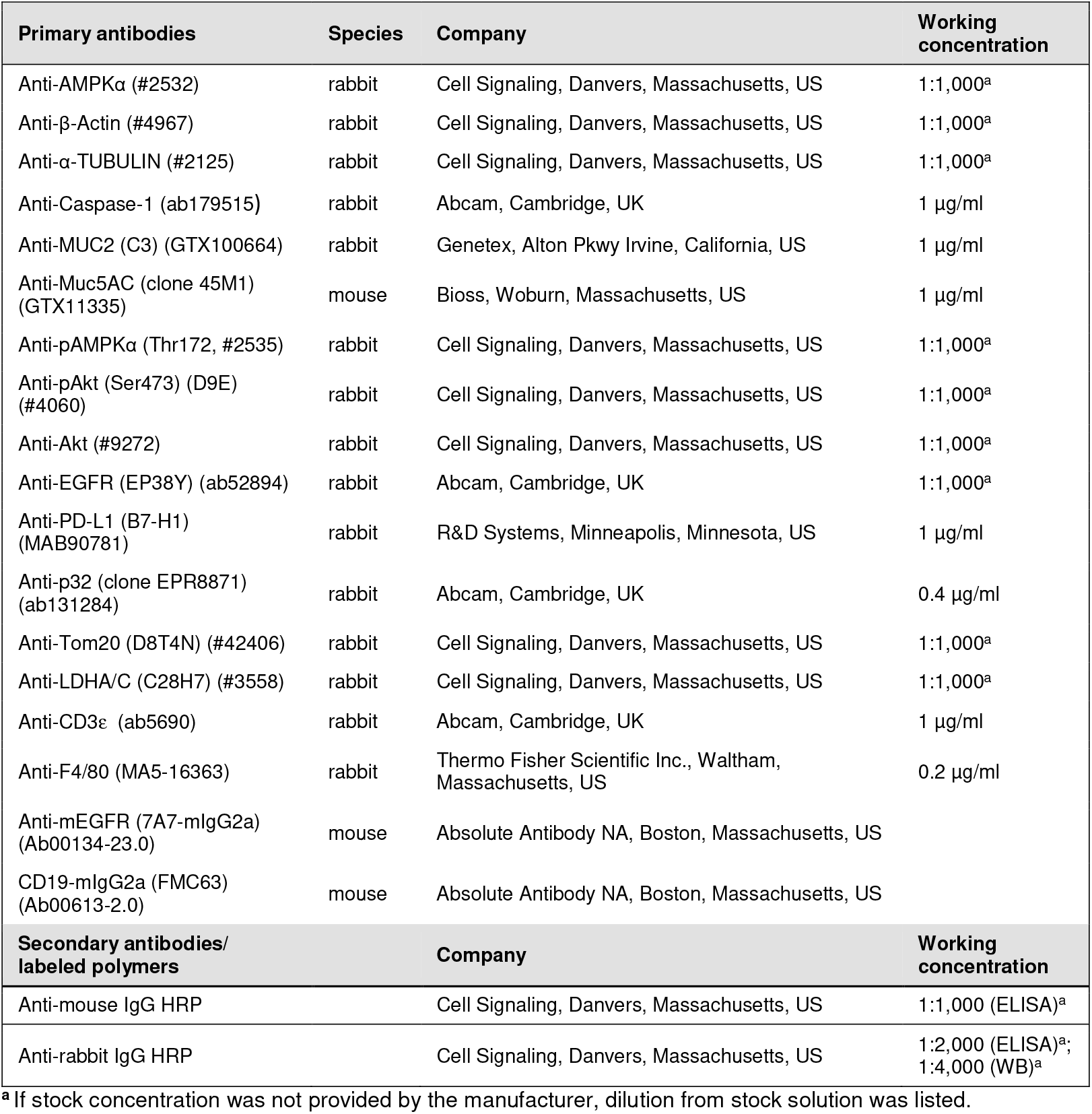
Details of applied primary and secondary antibodies.

| Primary antibodies | Species | Company | Working concentration |
| --- | --- | --- | --- |
| Anti-AMPKα (#2532) | rabbit | Cell Signaling, Danvers, Massachusetts, US | 1:1,000<sup>a</sup> |
| Anti-β-Actin (#4967) | rabbit | Cell Signaling, Danvers, Massachusetts, US | 1:1,000<sup>a</sup> |
| Anti-α-TUBULIN (#2125) | rabbit | Cell Signaling, Danvers, Massachusetts, US | 1:1,000<sup>a</sup> |
| Anti-Caspase-1 (ab179515) | rabbit | Abcam, Cambridge, UK | 1 μg/ml |
| Anti-MUC2 (C3) (GTX100664) | rabbit | Genetex, Alton Pkwy Irvine, California, US | 1 μg/ml |
| Anti-Muc5AC (clone 45M1) (GTX11335) | mouse | Bioss, Woburn, Massachusetts, US | 1 μg/ml |
| Anti-pAMPKα (Thr172, #2535) | rabbit | Cell Signaling, Danvers, Massachusetts, US | 1:1,000<sup>a</sup> |
| Anti-pAkt (Ser473) (D9E) (#4060) | rabbit | Cell Signaling, Danvers, Massachusetts, US | 1:1,000<sup>a</sup> |
| Anti-Akt (#9272) | rabbit | Cell Signaling, Danvers, Massachusetts, US | 1:1,000<sup>a</sup> |
| Anti-EGFR (EP38Y) (ab52894) | rabbit | Abcam, Cambridge, UK | 1:1,000<sup>a</sup> |
| Anti-PD-L1 (B7-H1) (MAB90781) | rabbit | R&D Systems, Minneapolis, Minnesota, US | 1 μg/ml |
| Anti-p32 (clone EPR8871) (ab131284) | rabbit | Abcam, Cambridge, UK | 0.4 μg/ml |
| Anti-Tom20 (D8T4N) (#42406) | rabbit | Cell Signaling, Danvers, Massachusetts, US | 1:1,000<sup>a</sup> |
| Anti-LDHA/C (C28H7) (#3558) | rabbit | Cell Signaling, Danvers, Massachusetts, US | 1:1,000<sup>a</sup> |
| Anti-CD3ε (ab5690) | rabbit | Abcam, Cambridge, UK | 1 μg/ml |
| Anti-F4/80 (MA5-16363) | rabbit | Thermo Fisher Scientific Inc., Waltham, Massachusetts, US | 0.2 μg/ml |
| Anti-mEGFR (7A7-mIgG2a) (Ab00134-23.0) | mouse | Absolute Antibody NA, Boston, Massachusetts, US |  |
| CD19-mIgG2a (FMC63) (Ab00613-2.0) | mouse | Absolute Antibody NA, Boston, Massachusetts, US |  |
| Secondary antibodies/<br>labeled polymers |  | Company | Working concentration |
| Anti-mouse IgG HRP |  | Cell Signaling, Danvers, Massachusetts, US | 1:1,000 (ELISA)<sup>a</sup> |
| Anti-rabbit IgG HRP |  | Cell Signaling, Danvers, Massachusetts, US | 1:2,000 (ELISA)<sup>a</sup>;<br>1:4,000 (WB)<sup>a</sup> |

2 <sup>a</sup> If stock concentration was not provided by the manufacturer, dilution from stock solution was listed.

### Neutral red cytotoxicity assay

The neutral red cytotoxicity assay was performed to determine the viable cell mass of MC-38 cells. 5 x 10^3^ cells per well were seeded into a 96-well microtiter plate and incubated at 37°C and 5% CO_2_ for 72 hours. After incubation, cells were stained using a neutral red dye (Sigma-Aldrich) diluted 1:100 in DMEM medium for 2 hours, washed and destained with a solution consisting of 50% pure ethanol, 49% bidistilled water and 1% pure acetic acid to release the incorporated dye into the supernatant. To analyze the neutral red dye uptake, absorbance was measured at 540 nm against a background absorbance of 690 nm on a SpectraMax iD3 microplate reader (Molecular Devices).

### Muc2 and Muc5AC ELISA

For detection of Muc2 or Muc5AC secretion by MC-38 or HT29-MTX cells, respectively, supernatants were centrifuged at 12,000 x *g* and 4°C for 15 minutes before coating into highly absorptive 96-well microtiter plates at 4°C overnight. After blocking, Muc2 or Muc5AC were detected using primary antibodies specific for Muc2 or Muc5AC in combination with respective HRP-conjugated secondary antibodies listed in **table 3**. Optical density was measured at 450 nm against a reference wavelength of 540 nm on a SpectraMax iD3 microplate reader (Molecular Devices).

## Author approval

All authors had access to the study data, reviewed and approved the final manuscript.

## Statistics

Statistical analysis was performed using the GraphPad Prism version 8.0.1 (San Diego, California, US). Outliers were identified by ROUT test. The F test was used to compare variances and D’Agostino– Pearson test was applied to test for normal distribution. Statistical differences between two groups were analyzed by unpaired t-test or paired t-test (normally distributed data), unpaired t-test with Welch’s correction (significant different variances) or Mann–Whitney U-test (not-normally distributed data). For comparison of more than two groups and two variables two-way analysis of variances (ANOVA) with uncorrected Fisher’s Least Significant Difference test was employed. Correlation analysis was performed by obtaining the Pearson’s correlation coefficient. P-values were calculated and null hypotheses were rejected when *p* ≤ 0.05.

## Abbreviations

(AMPK): Adenosine monophosphate-activated protein kinase
(AOM): azoxymethane
(ATOH1): atonal basic helix-loop-helix transcription factor 1
(BCMA): B-cell maturation antigen
(BCAA): branched-chain amino acids
(ctrl): control
(CD): control diet
(CRC): colorectal carcinoma
(CTLA4): cytotoxic T-lymphocyte-associated protein 4
(DSS): dextran sodium sulfate
(DAI): disease activity index
(EGFR): epidermal growth factor receptor
(ELISA): enzyme-linked immunosorbent assay
(FCS): fetal calf serum
(GFHPD): glucose-free, high-protein diet
(HES1): hairy and enhancer of split-1
(HE): hematoxylin-eosin
(*i.p*.): intraperitoneal
(KLF4): Kruppel-like factor 4
(LDHA/C): lactate dehydrogenase a/c
(LY6G): lymphocyte antigen 6 complex locus G6D
(mAb): monoclonal antibody
(LGR5): leucine-rich repeat-containing G-protein coupled receptor 5
(MUC2): mucin 2
(MUC5AC): mucin 5AC
(MEICS): murine endoscopic index of colitis severity
(NEAA): non-essential amino acids
(ND3): NADH dehydrogenase 3
(OXPHOS): oxidative phosphorylation
(PFA): paraformaldehyde
(PAS): periodic acid-Schiff
(PD-1): programmed cell death protein 1
(PD-L1): programmed death-ligand 1
(qPCR): quantitative polymerase chain reaction
(SPDEF1): SAM pointed domain-containing Ets transcription factor 1
(SDS-PAGE): sodium dodecyl sulfate polyacrylamide gel electrophoresis
(TOM20): translocase of the outer membrane 20 homolog
(TIL): tumor-infiltrating lymphocyte.

## Acknowledgements

The authors gratefully thank Dr. Nieves Baenas and Dr. Katharina Schlumm for helping with organ sampling during animal studies.

